# The effect of task demands on the neural patterns generated by novel instruction encoding

**DOI:** 10.1101/2021.03.08.434338

**Authors:** Alberto Sobrado, Ana F. Palenciano, Carlos González-García, María Ruz

## Abstract

Verbal instructions allow fast and optimal implementation of novel behaviors. Previous research has shown that different control-related variables structure neural activity in frontoparietal regions during the encoding of novel instructed tasks. However, it is uncertain whether different task goals modulate the organizing effect of these variables. In this study, we investigated whether the neural encoding of three task-relevant variables (dimension integration, response set complexity and target category) is modulated by implementation and memorization demands. To do so, we combined functional Magnetic Resonance Imaging (fMRI), an instruction-following paradigm and multivariate analyses. We addressed how and where distributed activity patterns encoded the instructions’ variables and the impact of the implementation and memorization demands on the fidelity of these representations. We further explored the nature of the neural code underpinning this process. Our results reveal, first, that the content of to-be-implemented and to-be-memorized instructions is represented in overlapping brain regions, flexibly using a common neural code across tasks. Importantly, they also suggest that preparing to implement the instructions increases the decodability of task-relevant information in frontoparietal areas, in comparison with memorization demands. Overall, our work emphasizes both similarities and differences in task coding under the two contextual demands. These findings qualify the previous understanding of novel instruction processing, suggesting that representing task attributes in a generalizable code, together with the increase in encoding fidelity induced by the implementation goals, could be key mechanisms for proactive control in novel scenarios.

## 1. INTRODUCTION

Instruction following allows humans to implement novel behaviors quickly without prior practice. Such enhanced efficiency frames this ability as a special instance of humans’ cognitive flexibility (Cole, Laurent, & Stocco, 2013). An important yet unanswered question is how the brain rapidly reformats and organizes the symbolic information conveyed by instructions into efficient action (Brass, Liefooghe, Braem, & De Houwer, 2017). Recent findings have shown that the preparation to execute a novel instruction entails structured activity patterns across fronto-parietal regions, where the information encoding is organized by relevant dimensions of the current instructed task (Palenciano, González-García, Arco, Pessoa, & Ruz, 2019). However, it remains unknown whether this mechanism is triggered by the preparation to implement the novel tasks, or alternatively, whether it reflects mere declarative maintenance of the instructed content.

Prior studies have reported the pervasive effects of instruction following on behavioral and neural markers. The intention to execute a recently encoded instruction induces brain activation in areas associated with control and category-selective perceptual processing (González-García, Arco, Palenciano, Ramírez, & Ruz, 2017; González-García, Formica, Wisniewski, & Brass, 2021) and impacts the neural patterns representing the instructed content (Bourguignon, Braem, Hartstra, De Houwer, & Brass, 2018; Muhle-Karbe, Duncan, De Baene, Mitchell, & Brass, 2017; Ruge, Schäfer, Zwosta, Mohr, & Wolfensteller, 2019). In this line, a recent study (Palenciano, González-García, Arco, Pessoa, et al., 2019) showed that, during implementation demands, neural patterns in areas such as the inferior frontal gyrus (IFG) are organized by the need to integrate information from different dimensions of the instruction (e.g. color and size of the stimuli), while patterns in the intraparietal sulcus (IPS) and pre-supplementary motor area (pre-SMA) represent the complexity of the newly instructed response sets (Palenciano, González-García, Arco, Pessoa, et al., 2019).

Another set of neuroimaging studies from the instruction following literature has focused on how the declarative information of the instruction is transformed into a procedural format, that is, into an action-oriented proactive binding of relevant motor and perceptual codes (see Brass et al. 2017 for a review). This transformation (often defined as proceduralization) enables a highly accessible task model containing the condition-action rules from the instruction ready to be implemented, creating an optimal preparatory and reflexive-like state that enables a fast and optimal execution (Brass et al., 2017; but see Liefooghe & De Houwer, 2017). Interestingly, brain areas linked to novel instruction processing (such as the dorsal premotor cortex and middle frontal gyrus) seem to be involved also when participants only need to declaratively memorize (but not execute) the content of the instruction. Critically, these areas show higher levels of activation or information decodability under implementation demands (Bourguignon et al., 2018; Muhle-Karbe et al., 2017). This raises the question of how exactly declarative and procedural neural states differ.

A prominent theoretical proposal of instruction processing puts forward a three-step model of how the information contained in verbal instructions is transformed into action plans (Brass et al., 2017). First, the instruction content has to be encoded, building the representation of the declarative information and rules that specify the proper response (Hartstra, Waszak, & Brass, 2012; Sakai, 2008). Afterward, a preparation stage takes place before response execution (Sakai, 2008), where the task set is assembled and its proactive maintenance induces an adjustment of the features relevant to achieve the task (González-García et al., 2017; Muhle-Karbe et al., 2017; Ruge, Jamadar, Zimmermann, & Karayanidis, 2013). Finally, the task-set is executed and the action conveyed by the instruction is carried out (Stocco, Lebiere, O’Reilly, & Anderson, 2012). As mentioned before, proactive control not only biases perceptual and motor systems to enhance processing of upcoming stimuli and relevant responses, but it also organizes neural activity according to control-related variables, such as the need to integrate dimensions, or response complexity (Cole, Patrick, Meiran, & Braver, 2018; Palenciano, González-García, Arco, & Ruz, 2019). However, it remains unknown whether the reported proactive pattern organization in control-related areas enables the preparatory state that ultimately leads to execution, or it underlies the declarative memorization of the novel task demands.

The current study aimed to test the extent to which proactive pattern organization in cognitive control-related, motor, and perceptual regions is specific to the proceduralization of novel instructions. To that end, we adapted an instruction-following fMRI paradigm (González-García et al., 2017; Palenciano, González-García, Arco, & Ruz, 2019; Palenciano, González-García, Arco, Pessoa, et al., 2019) in which novel verbal instructions had to be either implemented (proceduralized) or memorized (non-proceduralized). As in Palenciano et al. (2019), the instructions from both conditions were manipulated according to three variables related to proactive preparation: 1) the necessity to integrate within or across stimuli dimensions, 2) the response set complexity and 3) the relevant target category. Using multivariate pattern analysis (MVPA; Haxby, Connolly, & Guntupalli, 2014) we aimed to explore the strength with which control-related variables organize brain activity patterns during both implementation and memorization demands (Bourguignon et al., 2018; Muhle-Karbe et al., 2017; Palenciano, González-García, Arco, Pessoa, et al., 2019). We expected, first, that the instruction structure would be represented, during task encoding, in several areas in both implementation and memorization conditions (Muhle-Karbe et al., 2017). More specifically, and in line with previous findings (Palenciano, González-García, Arco, Pessoa, et al., 2019), we predicted that activity patterns in the IFG would encode the kind of dimension integration required by the instruction, in the pre-SMA and IPS, the response complexity, and in the ventral visual pathway, the relevant target category. Second, and more importantly, we hypothesized that the effect of these variables on activity patterns would be emphasized under implementation demands – reflecting their role during the proceduralization process. In consequence, we expected higher decoding accuracy of such variables in the implementation than in the memorization condition.

## 2. METHODS

### 2.1. Participants

Thirty-seven students from the University of Granada (Spain) took part in the study (29 females, 8 males, mean age = 22.97, SD = 3.42). The participants were native Spanish speakers, right-handed and with normal or corrected-to-normal vision. They received economic compensation (20-35€, depending on performance) for their participation. They all signed a consent form approved by the Ethics Committee for Human Research of the University of Granada. Two participants were excluded due to excessive head movement (> 3mm) and other three due to low performance (< 75% of correct responses), resulting in a final sample of 32 participants. The sample size (N=32) was calculated a priori with the power analysis software PANGEA (Westfall, 2015), to detect with 80% power a small-medium (*Cohen’s d* = 0.3) two-way interaction effect in our behavioral analyses (see section *3.1. Behavioral results*).

### 2.2. Stimuli, apparatus

The stimuli consisted of 192 different verbal instructions (taken from Palenciano, González-García, Arco, Pessoa, et al., 2019). They all had the same “If …. then …” structure and were composed of two conditional statements and two responses (e. g.: “*If there are two vegetables and one fruit, press A. If not, press L*”). “A” and “L” corresponded to a left- and a right-hand response, respectively. Participants used their middle fingers for one task (e.g., implementation) and the index fingers for other (e.g., memorization). The finger assigned to each task was counterbalanced across participants. All the instructions referred to different features of either human faces or food items. The faces’ features were their gender (male, female), emotion (happy, sad), race (black, white), and size (large, small). The food items’ features were their type (vegetable, fruit), color (green, yellow), shape (elongated, rounded), and size (large, small). Equivalent face-related and food-related instructions were created by equating the features gender and food type, emotion and color, and race and shape across the two categories. Given that targets consisted of a grid of 8 stimuli (see below), instructions also specified the number of items from the grid that had to be taken into consideration to respond (one, two, or three).

Critically, we manipulated the structure of the instructions across three variables. First, the dimension integration, with instructions requiring the integration of features from the same dimension (e.g. “If there are two women and one man”, the integration is *within* the gender dimension) or from different dimensions (“If there are two women and one happy person”, the integration is *between* the gender and the emotion dimensions). Second, we manipulated the response set complexity, as the responses required could be single (e.g., “press A. If not press L”) or sequential (e.g., “press AL. If not press LA”). And third, the relevant target category, with instructions referring either to faces or to food items. These three variables were orthogonally manipulated in a trial-by-trial fashion. For instance, the instruction “If there are two women and one happy person, press A. If not, press L.” would belong to the between dimension integration, single response set, face-related condition.

Additionally, we manipulated the task demands (implementation vs. memorization) between blocks, similar to previous studies (e.g., Muhle-Karbe et al., 2017). In implementation blocks, each instruction was associated with a grid of target stimuli that either fulfilled the relevant conditions (50% of trials) or not (50% of trials). All grids consisted of combinations of 4 faces and 4 food items, drawn from a pool of 16 stimuli: 8 faces (4 men, 4 women, 4 happy, 4 sad, 4 white people, 4 black people, extracted from the NimStim database; Tottenham et al., 2009) and 8 food items (4 vegetables, 4 fruits, 4 yellow, 4 green, 4 elongated, 4 rounded; extracted from Palenciano, González-García, Arco, Pessoa, et al., 2019). The 16 stimuli could appear in big or small size, generating a pool of 16 items per category (32 in total). All grids were created with a specific combination of stimuli, and each one appeared only once during the whole experiment.

Memorization blocks were identical to the implementation ones concerning the instructions. However, instead of grids of stimuli, targets consisted of another instruction that could be the same (50% of trials) or different (50% of trials) from the one encoded first. Different target instructions were created by exchanging either one of the stimulus features (e.g., “If there are two vegetables and one yellow food item” instead of “If there are two vegetables and one green food item”, where the feature color changed from yellow to green) or the response (e.g., “press A. If not press L” instead of “press LA. If not press AL”) from the original one. Target instructions also differed from the first encoded ones in font size and case (lower and uppercase for the encoded and target instruction, respectively). This manipulation sought to avoid that participants responded based on perceptual invariance or physical changes between the two instructions. In the target screen, the words “*different*” and “*same*” appeared at the right and the left side on the bottom of the screen, indicating the trial’s relevant mapping (e.g.: left middle finger if the instruction was the same, right middle finger if it was different). The response mapping changed across trials to prevent any potential response preparation before target onset as has been done in preceding studies (Formica, González-García, Senoussi, & Brass, 2021). In the scanner, the main task was organized in 12 blocks (6 implementation and 6 memorization ones). Implementation and memorization blocks were presented in alternation, with order counterbalanced across participants. Each block contained 16 trials (192 trials in total: 96 implementation and 96 memorization ones).

The task was presented through a screen connected to a computer running Matlab with the Psychophysics Toolbox (Brainard, 1997), with a set of mirrors mounted on the head coil, allowing the participants to see the screen. Responses were given through an MRI-compatible response pad with the middle and index fingers of both hands.

### 2.3. Procedure

The day before the scanning session, participants completed a practice where they were instructed about the tasks to be performed and repeated implementation and memorization blocks until they achieved 85% of accuracy. Participants completed at least two block repetitions per task demands were required. If they did not reach the established accuracy level after three practice repetitions, they were not invited to the scanning session (receiving an economic compensation for the time spent, at an hourly rate of 5 Euros). Six participants were not able to achieve 85% accuracy and therefore not further tested. All materials employed in the practice blocks were different from those used in the fMRI session.

In the scanner, at the beginning of each block, a verbal cue indicated the task to be performed (implementation vs. memorization). All trials (see Fig. 1) began with an instruction (*encoding stage*; 2.5s; 25.75°), which needed to be later executed (implementation) or remembered (memorization). Then, a fixation point (0.5°) was presented during a jittered interval ranging from 4 to 7.5s in steps of 500ms (average duration = 5.75s). This jitter was followed by a target grid (2.5s; 21°) in implementation trials, and by another instruction (2.5s; 28.5°) in memorization ones. Participants responded with the middle fingers for one task (e.g., implementation) and the index fingers for other (e.g., memorization), counterbalanced across runs. In both task demands, trials ended with a jittered fixation with the same characteristics as the previous one. In all cases, instructions were always new and were presented only once during the entire experiment.

**Figure 1.**
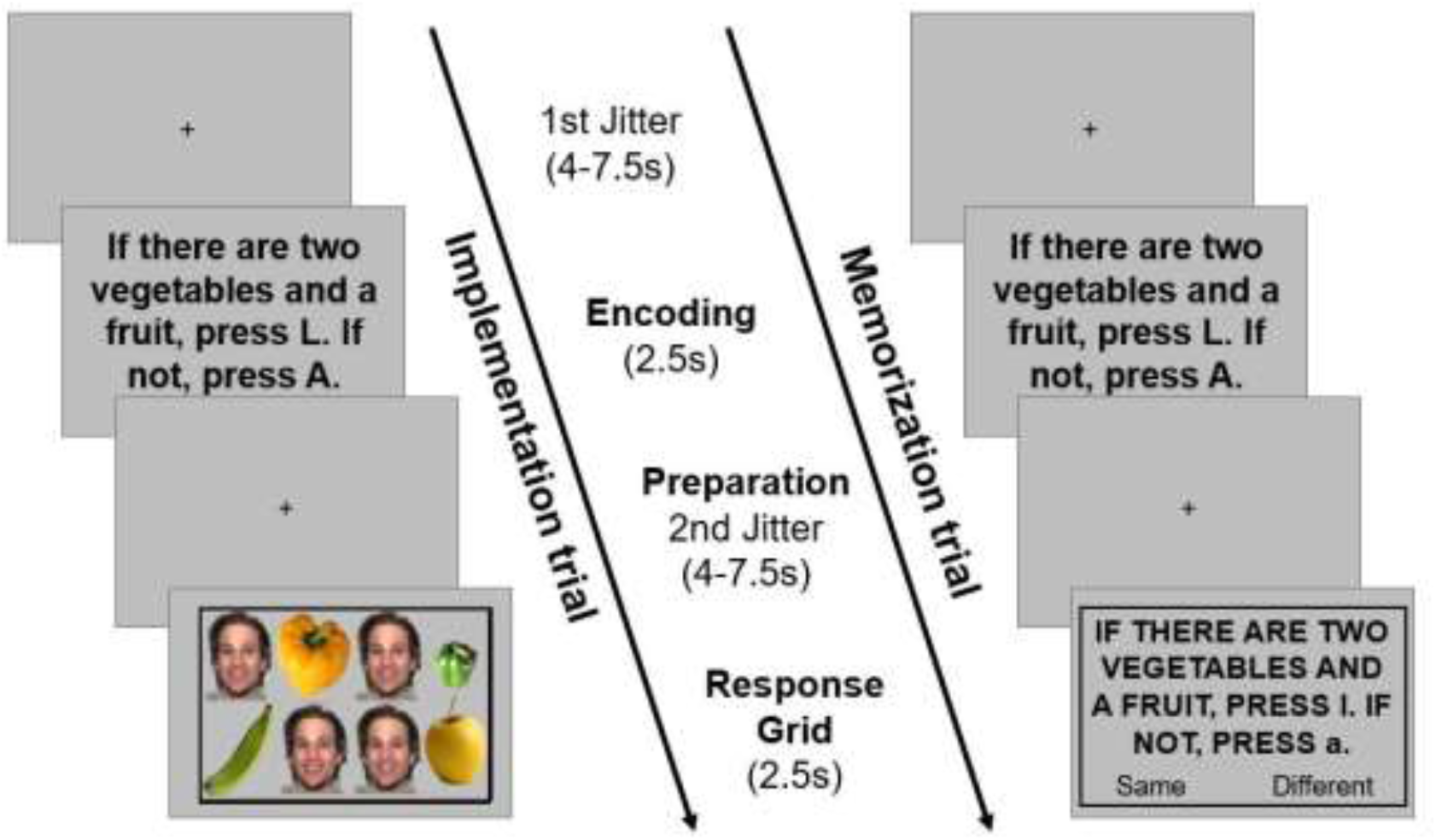
Behavioral paradigm.

Our four independent variables (task demands, dimension integration, response set complexity and target category) followed a within-subject design, with task demand being manipulated between blocks, and the remaining variables, randomized within-blocks. The distribution of jitters’ duration was equated across blocks. Participants spent ~90 minutes in the scanner, with the main task lasting ~75 minutes.

### 2.4. fMRI: acquisition and preprocessing

Participants’ MRI data were acquired with a 3T Siemens Magneton Trio scanner located at the Mind, Brain, and Behavior Research Center (University of Granada, Spain). Functional images were collected using a T2*-weighted echo-planar imaging (EPI) sequence (TR = 2000ms, TE = 24ms, flip angle = 70°). Each volume consisted of 34 slices, obtained in descending order, with 3.0 mm thickness (gap 20%, voxel size = 3 mm^3^). For each participant, a total of 1740 volumes were obtained, in 12 runs of 145 volumes each. Additionally, we acquired a structural image with a high-resolution anatomical T1-weighted sequence (192 slices of 1 mm, TR = 2500ms, TE = 3.69ms, flip angle = 7°, voxel size = 1mm^3^).

We used SPM12 to preprocess and analyze the data. The first 4 volumes of each run were discarded to allow stabilization of the signal. The remaining volumes were spatially realigned, unwarped and slice-time corrected. Then, the anatomical T1 was coregistered to the realigned functional images and segmented into different brain tissues. The deformation fields thus obtained were used to normalize the functional data to the MNI space (3mm^3^ voxel size). Last, the images were smoothed using an 8mm Gaussian kernel. Multivariate analyses (see below) were conducted with non-normalized and unsmoothed images, normalizing and smoothing the output maps before the group analyses.

### 2.5. fMRI: univariate analysis

For each participant, we estimated a General Linear Model (GLM) with separate regressors for each experimental condition (32 in total, resulting when crossing all the manipulation levels of our four independent variables). We defined two events per trial: the instruction and grid, both modeled with their duration (2.5s) and convolved with the canonical hemodynamic response function. Every regressor was modeled as a combination of the two trials of the same condition per run. Jitters were not modeled and thus contributed to the implicit baseline. As nuisance regressors, we included the 6 motion parameters and errors trials (modeled with the full duration of instruction, grid, and both jitters).

We focused on the trials’ encoding stage, to avoid potential confounds from perceptual (grid of stimuli vs. instruction) or motor differences (single or sequential responses vs. fixed single response) at the target stage between implementation and memorization conditions. At the individual level, we obtained maps contrasting the implementation and memorization demands during instruction encoding in both directions (implementation > memorization, and the other way around). These images were then entered into group-level one-sample *t-*tests, to identify brain regions with different mean activations depending on task demands. The results were corrected for multiple comparisons using an FWE cluster-level threshold of *p* < .05 (computed from an uncorrected *p* < .001).

### 2.6. fMRI: multivariate analyses

Our main goal was to explore how task demands modulated the encoding of novel instructions’ structure. First, we specifically addressed whether implementation demands entailed stronger, more distinguishable neural representations of the instructions variables manipulated in our design. Next, we further assessed if a generalizable neural code was shared by both task demands. Finally, we explored whether the encoding of to-be-implemented and to-be-memorized instructions also involved different, non-overlapping brain regions. It is important to stress that across all the analyses, we decoded the three instruction’s variables, instead of directly classifying between task demands (i.e.: between implementation and memorization trials). This approach ensured that our results were not contaminated by motor confounds derived from the different responses required by each task.

#### 2.6.1. Whole-brain decoding analyses and ROI selection

Our first goal was to detect brain areas coding each instruction’s variables (dimension integration, response set complexity, target category) during the encoding stage. To do so, we carried out three whole-brain MVPAs, one per manipulated variable. Since both memorization and implementation tasks were identical at the instruction encoding event, we collapsed across both demands (following a similar approach as in Palenciano et al., 2019). This way, we capitalized on the full dataset to maximize statistical power. From these results, we created Regions of Interest (ROIs) at the single-subject level, where we assessed the effect of task demands (implementation, memorization) on the fidelity of instructions’ variable encoding (see section *2.6.2. ROI-based decoding)*.

To carry out this and the remaining multivariate analyses, we estimated trial-by-trial GLMs with non-normalized and unsmoothed data. A Least-Square Separate approach (LSS; Arco, González-García, Díaz-Gutiérrez, Ramírez, & Ruz, 2018; Mumford, Davis, & Poldrack, 2014) was followed to gain sensitivity and to reduce collinearity between regressors. With the trial-wise beta coefficient maps from the encoding stage, we trained three Support Vector Machine classifiers to decode between the two levels of each of the three instructions’ variables. In the dimension integration MVPA, we classified between instructions integrating within and between dimensions. For the response set complexity, we decoded between instruction requiring simple and sequential responses. Finally, in the target category MVPA, we classified between faces and food-related trials. The classification was performed in every location of the brain, iterating a 4-voxel radius searchlight sphere. We used a 6–fold cross-validation scheme, training the classifiers in 10 out of the 12 runs of the experiment (five implementation and five memorization blocks) and testing them in the remaining two runs (one per task demand). We iterated this procedure until all runs were used as test data once. To avoid a biased classification performance, we ensured a balanced presence of implementation and memorization data across the cross-validation procedure. After averaging across cross-validation folds, the resulting subject-wise balanced accuracy maps were introduced into a group-level one-sample *t*-test against chance. This way, we obtained three group maps, each one displaying significant above-chance decoding of a particular instructions’ variable. The results were corrected for multiple comparisons with a cluster-wise FWE threshold at *p* < .05 (from an initial, uncorrected threshold at *p* < .001).

#### 2.6.2. ROI-based decoding

Next, we used as candidate ROIs the areas identified with the whole-brain classification, where we assessed the effect of the task demands on novel instruction encoding. In particular, we tested whether implementation demands increased the decodability of the three instructions’ variables. To avoid circularity in this analysis, the ROIs were derived for each participant using a Leave-On-Subject-Out (LOSO) approach (Esterman et al., 2010). We repeated the three group-level *t*-tests (decoding accuracy against chance for dimension integration, response set complexity, and target category) for each participant, excluding their own data from the analyses. The significant clusters from each LOSO map (thresholded at *p* < .001) that matched the areas resulting at the group level were used as ROIs for that particular participant. Since not all clusters obtained at the group level were present across all the LOSO estimations, we excluded from further analysis ROIs shared by less than 25 participants (i.e., 80% of our sample). The resulting subject-wise ROIs were inverse-normalized to the participants’ native space.

In each of these ROIs, we classified between the two levels of the corresponding instruction variable, separately for implementation and memorization data. As a result, we obtained a decoding accuracy value per participant, ROI and task demand. To compare the encoding strength between the implementation and the memorization tasks, we carried out paired-sample Wilcoxon signed-rank tests. As a sanity check, we used one-sample Wilcoxon signed-rank tests to assess above-chance decoding accuracy within each region. All results were corrected for multiple comparisons (i.e., the number of ROIs analyzed) using a *p* < .05 Bonferroni-Holm criteria (Holm, 1979).

#### 2.6.3. Whole-brain cross-classification

To have further insights on how novel instructions were encoded under different demands, we explored whether the same or different neural codes were recruited by implementation and memorization tasks. To do so, we performed a cross-classification analysis (Kaplan et al., 2015). We trained classifiers to decode the instructions’ variables with one particular task (e.g., implementation) and assessed its performance on the other (e.g., memorization). A successful classification would be interpreted as evidence of generalizability between demands, that is, a common neural code underpinning instruction representation.

The cross-decoding followed the same logic and parameters as the former whole-brain decoding. Using a searchlight procedure, we trained separate SVM classifiers to decode the three instructions’ variables, following a 6-fold cross-validation scheme. The algorithm was now trained on five runs of one task (e.g., implementation) and tested on one run of the other (e.g., memorization), until all the runs were used as test data once. The process was then repeated with the opposite cross-decoding direction (e.g.: training in memorization, testing in implementation). The resulting decoding accuracy maps were averaged across cross-validation folds and cross-decoding directions and introduced into group-level one-sample *t*-tests against chance. The results were corrected with a *p* < 0.05 cluster-wise FWE threshold (from an uncorrected *p* < .001).

To compare the cross-decoding output with the more general whole-brain decoding results (*section 2.6.1.*), we additionally carried out a conjunction analysis, statistically assessing the anatomical overlap between these two approaches for each instruction variable. To do so, we computed the intersection of the thresholded group maps (cluster-wise FWE threshold at *p* < .05) obtained from each variable’s cross-classification and whole-brain decoding.

#### 2.6.4. Whole-brain within-task decoding

The former analyses addressed differences between implementation and memorization tasks on regions encoding instructions under both demands and which shared a common neural code. However, it could also be the case that either the neural populations or their underlying code were not shared by the two task demands. To explore this possibility, we carried out a final set of whole-brain MVPAs, decoding the three instructions’ variables separately for the implementation and the memorization task. The resulting accuracy maps (one per instruction variable and task) were contrasted against chance using one-sample *t*-tests. Finally, we carried out paired-sample *t*-tests contrasting the implementation and memorization maps to detect regions differentially engaged by each demand.

## 3. RESULTS

### 3.1. Behavioral results

Accuracy and reaction times (RT) were analyzed separately with repeated-measures ANOVAs using task demands (implementation, memorization), dimension integration (within, between dimensions), response set complexity (simple, sequential) and target category (faces, food items) as within-subject factors (main effects and interaction terms of interest for both ANOVAs are shown in Suppl. Table 1).

RT results evidenced the modulatory influence of the task demands over the effect of the remaining instructions’ variables. Specifically, task demands interacted significantly with the dimension integration, *F*(1,31) = 36.23, *p* < .001, *η*_*p*_^2^ = .54, response set complexity, *F*(1,31) = 17.02, *p* < .001, *η*_*p*_^2^ = .35, and target category, *F*(1,31) = 29.70, *p* < .001, *η*_*p*_^2^ = .49. We carried out post-hoc paired-sample *t*-tests to further describe these interactive patterns (Fig. 2). Regarding the dimension integration, participants responded generally slower when integrating between than within dimensions. The significant interaction showed that this effect was magnified under implementation, *t*(31) = 11.42, *p* < .001, than memorization demands, *t*(31) = 3.00, *p* = .004. Regarding the interaction of task demands with response set complexity, we found slower responses to sequential than single responses in the memorization task, *t*(31) = 3.40, *p* = .002, while smaller differences – that went in the opposite direction – were found in the implementation task, *t*(31) = −2.47, *p* = .016. Finally, regarding target category, participants were faster to food-related than to faces-related instructions in the implementation blocks, *t*(31) = 5.86, *p* < .001. This effect was not found in the memorization task *t*(31) = −1.09, *p* = .281.

**Figure 2.**
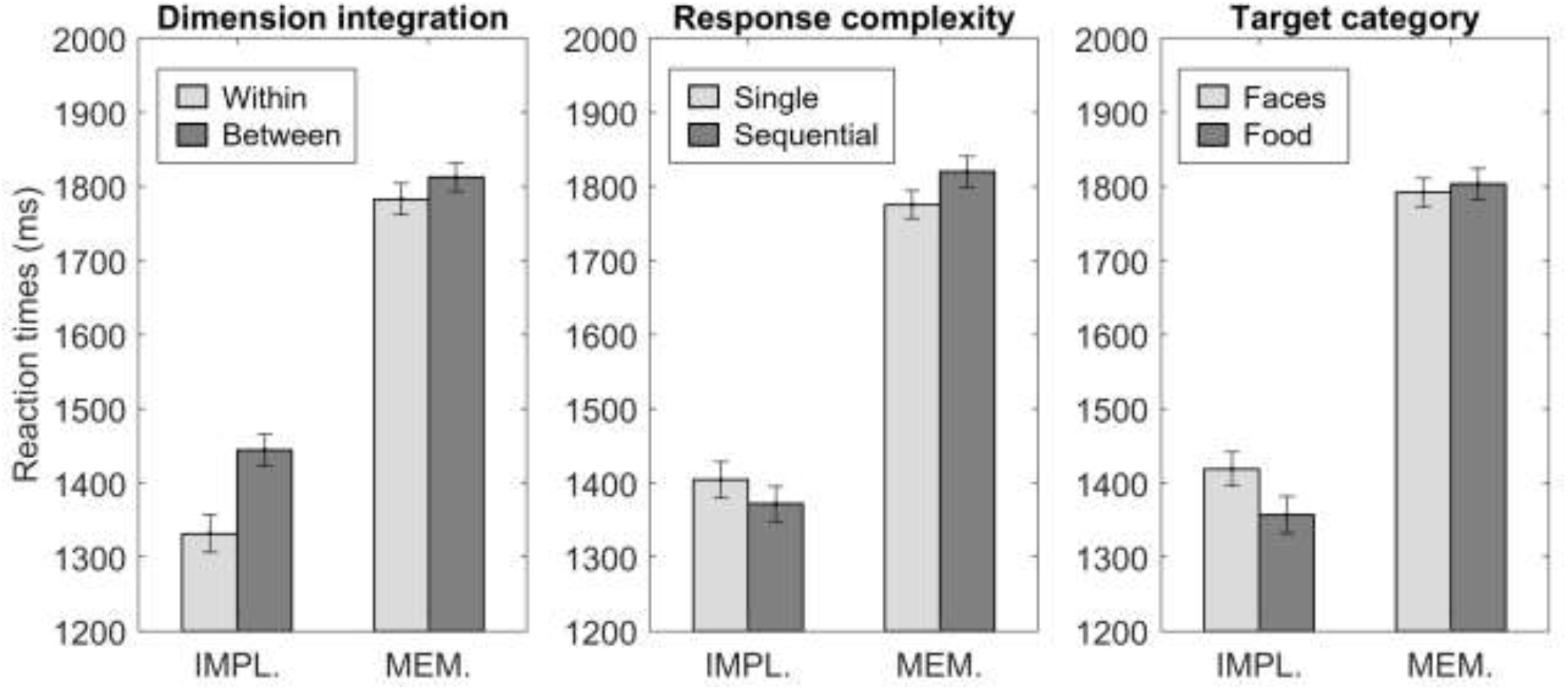
Mean RT for the two conditions of each instructions’ variable, separately for the implementation and the memorization tasks. Error bars depict standard errors. IMPL.: Implementation; MEM.: Memorization; Within: integration within dimensions; Between: integration between dimensions; Single: single response; Sequential: sequential response.

It is worth mentioning that responses were faster but also less accurate in implementation than in memorization trials, which could reflect a speed-accuracy trade-off induced by our task demand manipulation. To explore this, we performed a control analysis correlating differences in accuracy and RT between task demands across our sample. A significant, negative correlation would indicate that participants responding faster when implementing instructions would do so at the cost of more errors. Nonetheless, this analysis showed a negative but non-significant correlation (*r* = −0.257, *p* = .15), arguing against a trade-off interpretation.

### 3.2. fMRI: univariate results

First, we aimed to identify mean activation differences during instruction encoding under implementation and memorization demands. The brain maps showing the results for both tasks are displayed in Fig.3. The implementation demands induced greater activity than memorization ones in the bilateral fusiform gyrus (left: [−33, −37, −31], *k*=224; right: [33, −37, −31], *k* = 190), postcentral gyrus incurring into supramarginal and Rolandic operculum (right: [48, −19, 17], *k* = 97; left: [−54, −22, 14], *k* = 86), right supplementary motor area (SMA; [6, −4, 50], *k*=96), thalamus ([−3, −22, 11], *k* = 187) and right primary motor cortex (M1; [36, −13, 62], *k*=186). The memorization demands led to greater activity in comparison with the implementation condition only in the right angular gyrus ([24, −58, 44], *k* = 225).

**Figure 3.**
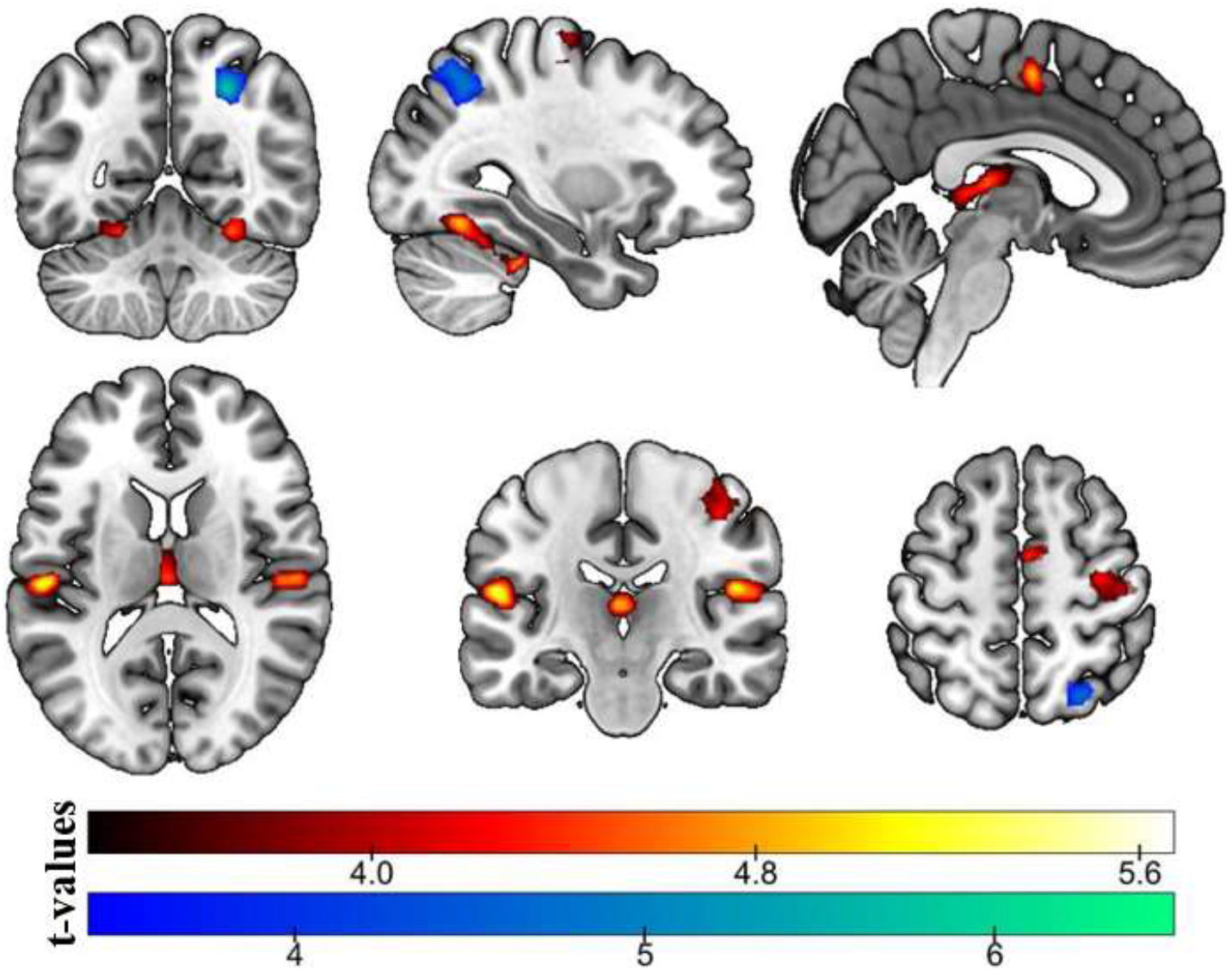
Univariate results. Regions more activated in implementation than memorization trials are shown in red, and the opposite contrast, in blue. The color shade indicates the *t*-values (*p* < .05, FWE corrected for multiple comparisons).

### 3.3. fMRI: Multivariate results

#### 3.3.1. Whole-brain decoding

As a first approach, we aimed to identify brain regions encoding the instructions variables, irrespectively of the task demand. Fig. 4.A shows the above-chance decoding accuracy maps for the three instructions’ variables. The dimension integration variable was successfully decoded only in the left middle temporal gyrus (MTG; [−57, −58, −7], *k* = 234). A less strict statistical threshold (*p*<0.001, uncorrected, see Suppl. Fig. 1) revealed an additional cluster encoding this variable in the left inferior frontal gyrus (IFG; [−57, 23, 17], *k* = 112), following our previous results (Palenciano, González-García, Arco, Pessoa, et al., 2019). The response complexity could be decoded from a wide cluster ([−45, −70, −1], *k* = 4594) spanning the bilateral premotor cortices (PMC), supplementary motor areas (SMA), and pre-SMA, and the left M1. The same cluster also extended posteriorly into the left somatosensory cortex and inferior parietal lobe (IPL), and into the left MTG and fusiform gyrus. Anteriorly, it also covered the left IFG and frontal operculum. Significant above-chance decoding of the response complexity was also found in the left middle frontal gyrus (MFG [-27, 56, 23], *k* = 175), and the right supramarginal gyrus (SMG, [54, −28, 38], *k* = 170). Finally, the target category was decoded from a large cluster in the left hemisphere ([−42, −43, −25], *k* = 2069) covering the left MTG and the inferior temporal gyrus, the fusiform gyrus, and the precuneus. The category was also decoded from the left IFG, extending into the frontal operculum and anterior insula ([−51, 32, 14], *k* = 1229), the dorsomedial prefrontal cortex (dmPFC; [−6, 53, 32], *k* = 455), and the ventromedial prefrontal cortex (vmPFC; [−3, 50, −22], *k* = 149).

**Figure 4.**
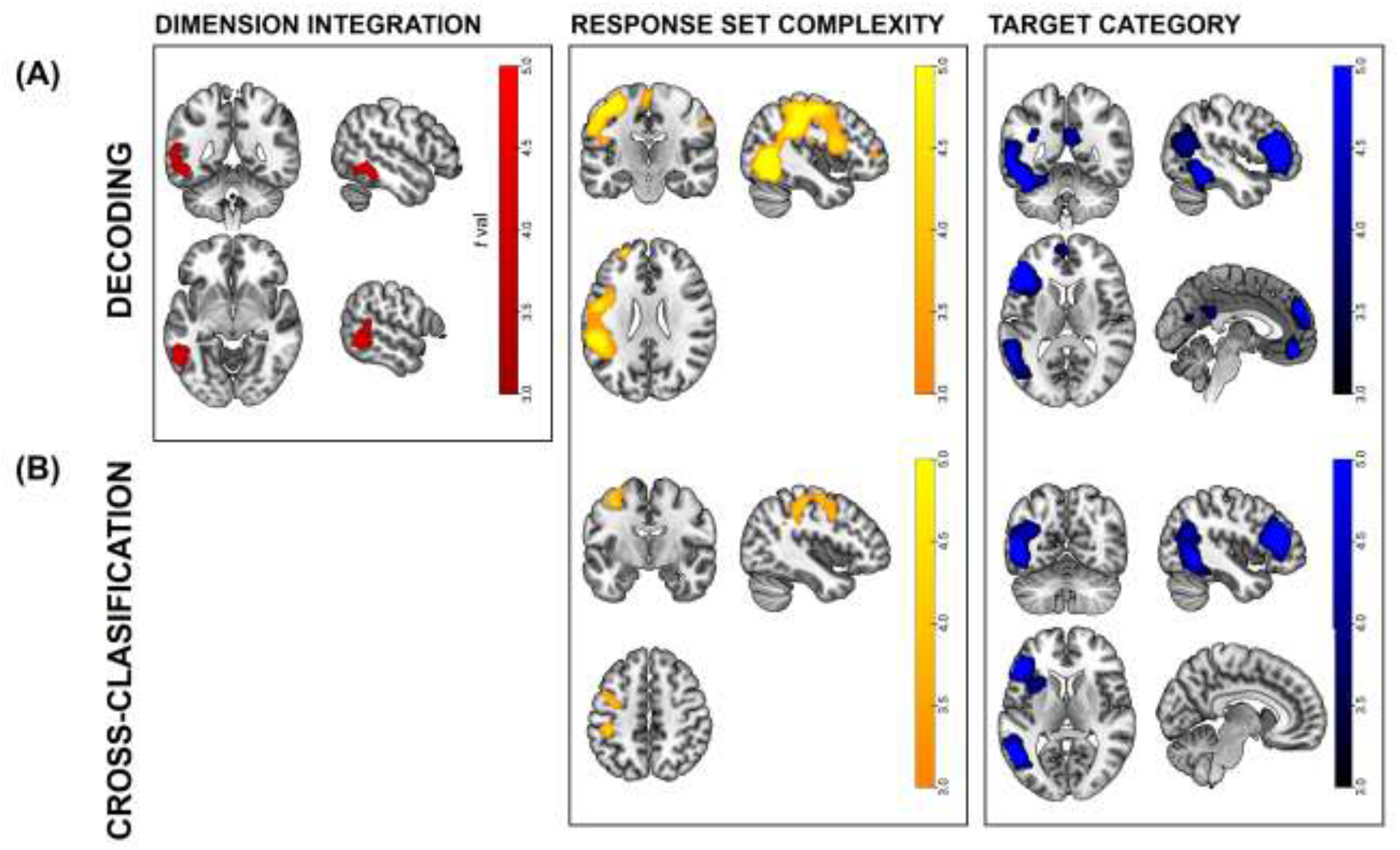
**(A)** Results from the whole-brain decoding analysis for each instructions’ variable. **(B)** Results from the whole-brain cross-classification analysis for each instructions’ variable. Color shades indicate *t*-values (*p* < .05, FWE corrected for multiple comparisons).

#### 3.3.2. ROI-based decoding

From the previous analyses, we extracted a series of ROIs encoding the instructions’ structure. Next, we investigated whether the intention to implement an instruction increased the fidelity (i.e., the decoding accuracy) of these representations, in comparison with memorization demands. After performing a LOSO procedure, we ended up with five ROIs: a left MTG ROI identified with the dimension integration decoding, an ROI extracted from the response complexity decoding which covered the motor cortices together with the IPL and left MTG, and three ROIs isolated with the target category decoding and located in the left MTG, left IFG and dmPFC. In each of these regions, we computed the classification accuracy for the corresponding variable, separately for implementation and memorization trials, and compared between task demands. Mean decoding accuracies per ROI and task are shown in Fig. 5. In line with our hypothesis, we found higher decoding accuracies under implementation demands for the response complexity variable across the motor cortices, the IPL and the left MTG, and for the target category variable, in the left MTG and IFG (see Table 1). No evidence in favor of this pattern (nor the opposite) was found regarding the integration of dimensions variable in the left MTG, nor the target category decoding in the dmPFC.

**Figure 5.**
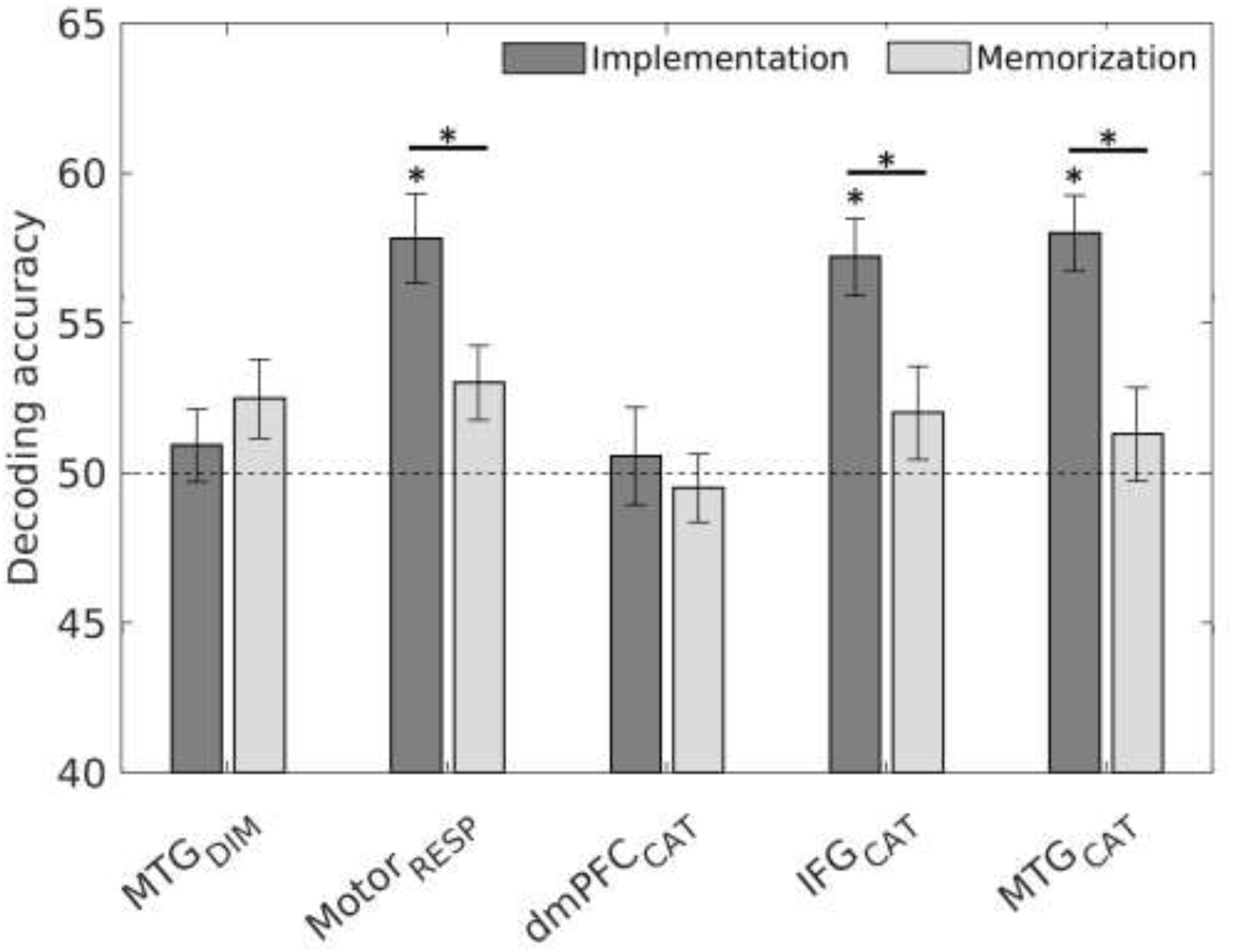
Mean decoding accuracy for each ROI and instruction’ variable, separately for the implementation and memorization conditions. The black dashed line shows chance decoding performance. Asterisks indicate significant results in the corresponding one-sample or paired-sample Wilcoxon sign rank test. Results are Bonforroni-Holm corrected for multiple comparisons. Error bars depict standard error. DIM: ROI extracted from the dimension integration decoding; RESP: ROI extracted from the response complexity decoding; CAT: ROI extracted from the target category decoding.

**Table 1.**
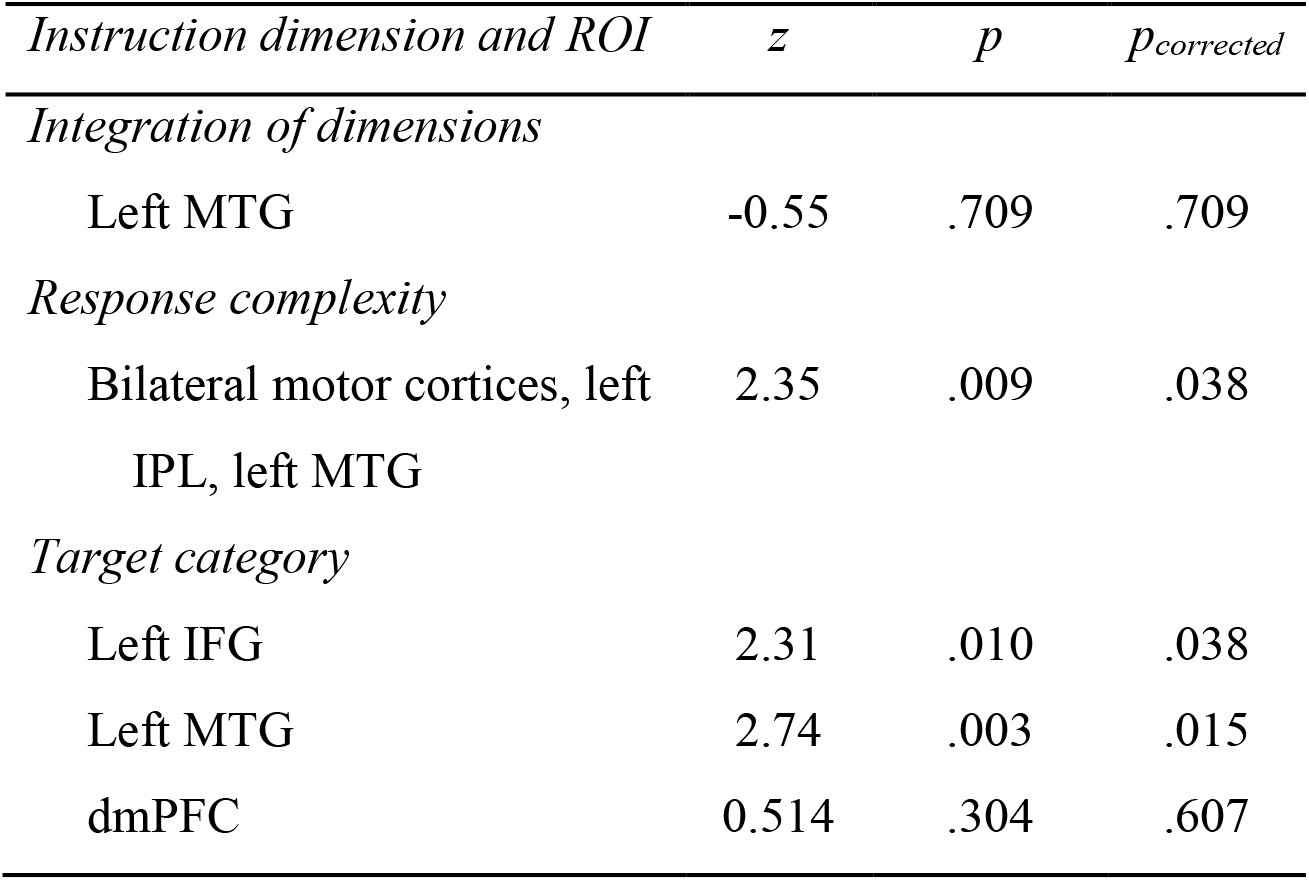
Results of the paired-sample Wilcoxon sign-rank test contrasting the decoding accuracy between implementation and memorization conditions

As a sanity check, we also tested whether the decodability found in these regions during the ROI-based analyses was significantly at chance (see Fig. 5). Critically, we did not find evidence supporting successful decoding of the dimension integration condition in the left MTG under neither demands. The same applied to the decoding of the target category in the dmPFC.

#### 3.3.3. Cross-classification

We further explored whether the neural code used to represent the different instructions’ variables was shared across task demands. To address that issue, we performed a cross-decoding analysis, testing whether a classifier trained in one task demand performed above chance in data from the other task. The brain maps from Fig. 4.B show significant cross-classification performance (the two cross-classification directions’ results are displayed in Suppl. Fig. 2). The response set complexity could be cross-decoded from the left PMC, M1 and IPL ([−34, −14, 68], k = 2134). In the target category analysis, we found significant clusters on the left IFG, including part of the frontal operculum and anterior insula ([−40, 38, 20], k = 3791) and the left MTG and fusiform gyrus ([−46, −56, −6], k = 3599). However, no evidence for above-chance cross-classification was found for the dimension integration.

To estimate the prevalence of common neural patterns between task demands, we carried out a conjunction analysis using the maps from the cross-classification and the whole-brain decoding analyses. Then, we computed the percentage of voxels identified with these two analyses which fell into the conjunction maps. Almost all voxels identified with the cross-classification analyses were captured by the conjunction (91.6% in the case of the response set complexity, and 80.1% for the target category), illustrating that the shared neural code took place majorly in regions already identified as encoding the instructions’ variables. In the case of the general decoding analyses, we found a 45.9% of overlap in the response set complexity classification, showing that the neural code generalized across demands in approximately half of the identified voxels. For the target category, we found a 12.1% of overlap between the general decoding and the conjunction maps, showing a lower but still considerable prevalence of shared neural code.

#### 3.3.4. Whole-brain within-task decoding

To extend the ROI-based results, we ran further whole-brain classification analyses separately for each task demand. We followed this approach to identify regions encoding novel instructions exclusively in implementation or memorization trials. This analysis also allowed us to detect differences in regions where the neural code was not common across demands (i.e., not captured by the cross-classification). The accuracy maps showing significant decoding of the three instruction variables for each task are displayed in Fig.6. While the necessity to integrate within or between dimensions was encoded in the left IFG ([−60, 17, 14], k = 168) in the implementation blocks, no significant clusters were detected in the memorization ones. Regarding response set complexity, we found significant above-chance decoding during the implementations blocks in the bilateral M1, PMC, and SMA extending into the bilateral intraparietal sulcus and IPL (left: [−45, −31, 44], k = 1424; right: [42, −34, 56], k = 783), in the left IFG and anterior insula ([−57, 5, 11], k = 179), and the left MTG ([−54,−61,−10], k = 270). The same classification in the memorization task showed significant clusters in the left IFG ([−54, 8, 17], k = 478), medial frontal cortex ([15, 38, −10], k = 399), and right cerebellum ([36, −76, −31], k = 345). Finally, during implementation blocks, the target category was encoded in the left IFG extending into the MFG ([−42, 29, 11], k = 796), the left fusiform gyrus ([−54, −52, −10], k = 533), and the precuneus ([−27, −58, 26], k = 630). No significant above-chance category decoding was found in the memorization blocks. Finally, we contrasted the implementation and memorization accuracy maps for each instruction variable. Nonetheless, no voxels survived in none of the comparisons.

**Figure 6.**
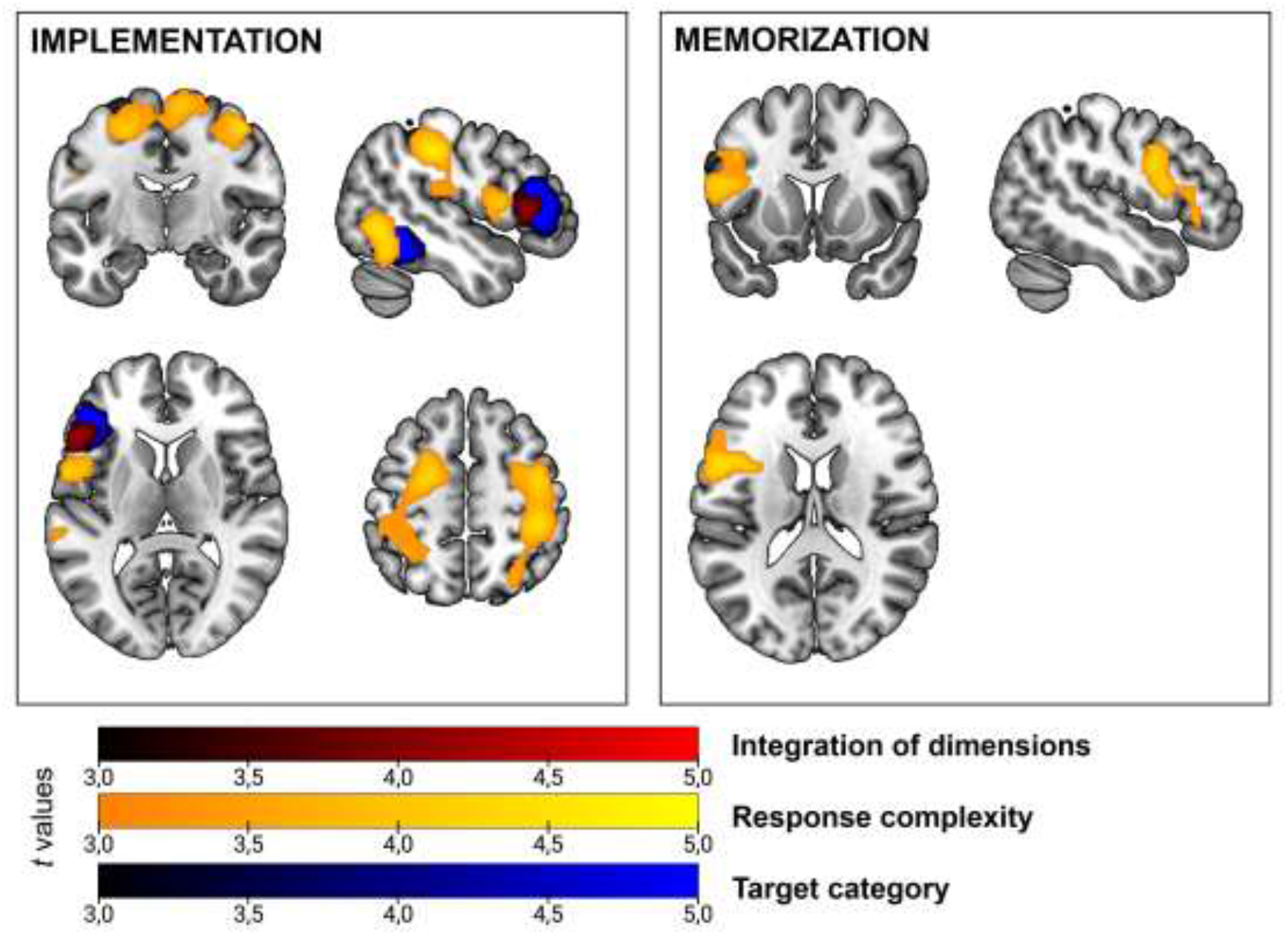
Results of the whole-brain within-task classification analyses. Color shades indicate *t*-values (p<.05, FWE corrected for multiple comparisons).

## 4. DISCUSSION

In the present study, we examined how task goals impact the structure of the neural patterns associated with the encoding of complex, novel instructions. Using multivariate analyses, we explored brain regions encoding instructions’ dimensions related to anticipatory control. Critically, we assessed whether the fidelity of the neural representations held in these areas was modulated by specific implementation or memorization demands. In addition, with cross-classification analysis, we aimed to deepen the understanding of the neural code supporting novel instruction processing across task demands. Finally, further within-task classification analyses explored whether the representation of the instructions’ dimensions could also take place on distinct neural populations. Our findings emphasize that both implementation and memorization demands share similar neural mechanisms during novel instruction encoding. Fronto-parietal, motor and perceptual regions represented the novel task content across both conditions, using a generalizable neural code. Nonetheless, our results also stress that to-be-implemented instructions induced more decodable representations of the task’s relevant variables, in comparison to memorization trials. This finding was further supported by the within-task classification results. Taken together, our work reveals a complex picture where cognitive control and sensorimotor areas process novel instructed content with a shared format for implementation and memorization tasks, but with different coding strengths.

The behavioral data already evidenced the impact of implementation and memorization demands on instructed performance. The participants’ response speed was sensitive to the instructions’ variables manipulated in our design (dimension integration, response set complexity, and task category), related to proactive task reconfiguration. Critically, the three variables’ effect was modulated by task demands. Integrating stimuli features between dimensions had a cost on performance in comparison with within-dimension integration. In line with our predictions, this cost was larger under implementation demands. Similarly, the target category had a larger effect on RTs in the implementation than in the memorization task, where no differences were found between faces and food-related instructions. This pattern may reflect the involvement of category-specific preparatory mechanisms on implementation trials, which would not be required when instructions were merely remembered. Finally, the response set complexity also interacted with task demands, this time with a greater effect under memorization demands. While longer RTs for sequential than single responses were found for the memorization task, an opposite, smaller effect was found in implementation demands. While this pattern deviated from our prediction, it converges with the remaining behavioral results, supporting that the variables manipulated were behaviorally relevant for instructed performance and showed specificity regarding task demands.

The univariate fMRI results showed that the encoding of to-be-implemented and to-be-memorized instructions increased the activity in different sets of brain regions, in line with previous research (Bourguignon et al., 2018; Muhle-Karbe et al., 2017). Replicating findings from past studies (e.g., González-García et al., 2017; Palenciano, González-García, Arco, & Ruz, 2019), the implementation condition entailed greater pre-activation during the encoding stage in areas associated with the proactive control-related instructions’ variables, as target category (fusiform gyrus) and response preparation (along the motor cortices). This increased activity may mediate the more effortful binding of perceptual and motor task information during the assembly of the procedural task representations. Interestingly, we also found higher activity in the thalamus for implementation than memorization blocks. This finding could reflect the need to integrate across multiple types of information (motor, perceptual, and abstract) and the maintenance of task representations (Wolff & Vann, 2019). Contrary, when instructions were encoded under memorization demands, higher activity was found in the angular gyrus. This area could participate in the elaboration of declarative instructions representations. Although this step is assumed to take place for both implementation and memorization demands (Brass et al., 2017), the declarative format would suffice to guide performance in the latter condition. This interpretation would be in line with previous research stressing the general role of the angular gyrus in declarative memory (Noonan et al., 2013; Seghier, 2013), and more in particular, with its involvement during the retrieval of rule-based schemas and their constituting components (Wagner et al., 2015). Altogether, our results could tentatively reflect a complex processing chain, in which the instructed task sets are built from its simpler constituents (Deraeve et al., 2019; Reverberi et al., 2012), first giving rise to a declarative memory trace, and if the current demands require it, transforming it into a procedural code (Brass et al., 2017; Cole et al., 2013).

Beyond activity mean differences, our main goal was to explore the fine-grained instruction structure coded in distributed activity patterns. Our findings replicated previous literature (González-García et al., 2017; Muhle-Karbe et al., 2017; Palenciano, González-García, Arco, Pessoa, et al., 2019), showing how a wide set of frontoparietal and sensoriomotor regions encode different task-relevant attributes in an anticipatory fashion. Critically, our work extended these results, directly investigating how these highly organized activity patterns are modulated by task demands. In this sense, our cross-classification analysis showed that prefrontal, motor and ventral visual cortices use a common neural code (Kaplan et al., 2015) to flexibly represent the instructions’ dimensions across tasks. In particular, significant generalization was found for the response set complexity and target category variables. That was not the case for the dimension integration variable - although the inference approach followed here does not enable the extraction of solid conclusions from these null results. Having said that, we can confirm the presence of a shared neural code at least regarding target stimuli and response-related information. These commonalities between the two task demands have been suggested in a previous study, which showed that the representations of to-be-implemented and to-be-memorized instructions are maximally similar during the encoding stage, when the decodability of task information is also equivalent between demands (Muhle-Karbe et al., 2017). Importantly, we further qualified these results, evidencing for the first time a common coding scheme in which novel instructions are represented irrespectively of the relevant demands. Moreover, this across-context generalizability has been proposed as an index of the abstraction achieved at representing task variables (Bernardi et al., 2020). Thus, the employment of abstract, shared task representations may be a key aspect for our flexible adaptation to novel demands (Badre et al., 2021; Bernardi et al., 2020). This interpretation also resonates with literature stressing the presence of compositionality underpinning complex task coding (Deraeve et al., 2019; Reverberi et al., 2012).

Critically, we also evidenced differences induced by implementation and memorization demands. We showed that the implementation task led to stronger, more distinguishable representations of target and response information. In our ROI analyses, we found this effect in three out of the five regions examined. In a similar vein, the within-task classifications showed that the three instructions’ variables could be decoded from a wide set of frontoparietal regions when only data from the implementation blocks were used, but not with the memorization data. While this set of results was not conclusive about differences between the two task demands (since no significant clusters were found when the two conditions were directly contrasted), these results converge with the ROI analyses’ ones. This general pattern, which goes in line with previous results (Muhle-Karbe et al., 2017), partially supports the hypothesis that implementation demands induce a boost in representational fidelity, which could potentially trigger the preparatory stage leading to efficient novel performance. As an alternative interpretation, difficulty differences between task demands could have also played a role in this pattern. In line with previous evidence (e.g., Woolgar et al., 2011), a more difficult implementation task could induce more decodable patterns in prefrontal and parietal regions. However, although our behavioral results showed more erroneous performance in the implementation task, they also evidenced slower response in memorization conditions, leaving unclear which demand was more difficult for our participants. Future studies explicitly manipulating task difficulty would be enlightening in this regard.

Overall, our findings point towards both similarities and distinctions during novel instruction coding depending on task goals. These results can shed some light on how instructions’ representations are quickly shifted from a declarative to an action-based or procedural format (Demanet et al., 2016; Hartstra et al., 2011; Muhle-Karbe et al., 2017). With our paradigm, we aimed to tap into these two stages, emphasizing the declarative phase in the memorization condition, and promoting the procedural reformatting under implementation demands. This way, we aimed to obtain new insights on how this transformation takes place. Past studies have proposed that the two stages are driven by distinguishable neurocognitive states, which in turn are reflected in representations of different nature (González-García et al., 2021; Muhle-Karbe et al., 2017). For instance, it has been recently evidenced that non-overlapping declarative and procedural formats are represented in fronto-parietal activity patterns (González-García et al., 2021). Our results, highlight a different, not mutually exclusive pattern: a continuity between declarative and procedural representations, with implementation goals amplifying or intensifying the instructed content. While ours and previous findings seem in contradiction, they could complement each other. For instance, it could be the case that the implementation-related boost in encoding strength (captured with our experiment) corresponds to an initial stage of the proceduralization of novel instructions. Later, it could lead to the clearer differentiation between procedural and declarative task representations found in previous studies. Future investigation using time-sensitive measures (as electroencephalografy recordings) woud be crucial to reconcile these two sets of results.

Even when the findings regarding the target category and response complexity variables were, in general terms, in line with our expectations, that was not the case for the dimension integration manipulation. Based on a previous study (Palenciano, González-García, Arco, Pessoa, et al., 2019), we predicted that neural patterns in the IFG would code the integration of information within or between stimulus dimensions, with greater decoding accuracies for implemented instructions. Instead, this region was not identified with our general decoding analysis. One potential explanation relates to our paradigm, which may have not generated enough separation between the two dimension integration conditions to engage the IFG. Nonetheless, and against that possibility, our manipulation replicated the one employed in Palenciano and collaborators (2019). Critically, when only data from the implementation task were used, this variable was indeed encoded in the left IFG. This result suggests that the processes carried out in this prefrontal region are of special relevance during the reformatting of the instruction content for the implementation. However, not enough evidence was obtained supporting specifically the link between the dimension integration encoding on the IFG and implementation demands. While a lack of statistical power may be the source of such null results, we still encourage caution when interpreting this particular finding, which should be confirmed in future research. Instead, our results highlighted the role of the MTG in encoding the dimension integration irrespectively of task demands. Interestingly, prior studies focusing on semantic retrieval had described a relationship between the left IFG and the MTG, which work coordinately in semantic demanding tasks (Davey et al., 2016). Taking that into account, our results could reflect the role of the MTG in the controlled retrieval of semantic information and its integration to correctly perform the action specified by the instruction.

It is important to stress that, when taking together the findings from the different analyses performed, we obtained a mixed pattern of results that requires further attention. On one hand, some of the regions encoding the instructions’ dimensions across task demands, identified with whole-brain decodings, were not significant in the ROI analysis. On the other, despite the within-task classification yielded different encoding maps when exploring the implementation and the memorization tasks separately, no differences were detected between both of them. This puzzling scenario could be the result of a lack of statistical power. In this sense, training our classifiers after splitting our data into implementation and memorization blocks (as it happened in the ROI and the whole-brain within-task analyses) could have hindered our ability to successfully decode the instructions’ content. That would especially affect the memorization condition, where smaller, more subtle effects would require more statistical power to be detected. Moreover, in the within task classification, a non-optimal statistical power could have interacted with the more stringent statistical correction required for whole-brain analyses, affecting the detection of significant differences between tasks. While we acknowledge this potential flaw, we encourage future, collaborative research initiatives which would provide wider datasets in which our main findings can be further replicated.

It is also worth noting that in this study, both univariate and multivariate analyses were performed at the encoding stage, while the instructions were displayed on the screen. That approach ensured avoiding possible confounds yielded by perceptual and motor differences during the response stage between the two task demands. This could be one key point in explaining some of the observed differences with prior studies characterizing the procedural and declarative processing of instructions. Most of this research focused on the preparation interval between the instruction coding and the motor response (Bourguignon et al., 2018; González-García et al., 2017; Muhle-Karbe et al., 2017), where the differences between both tasks are expected to be maximal (Brass et al., 2017; Liefooghe et al., 2013). The nature of our paradigm, however, did not allow us to focus our analyses on this preparation stage. Follow-up studies with optimized designs to extrapolate our findings to the preparatory stage would provide further information on this issue.

In sum, the present work examined the impact of implementation and memorization demands on new instructions processing, and in particular, how these two task goals modulated the neural patterns encoding several task-relevant dimensions. We show for the first time how frontoparietal, motor and perceptual brain regions represented novel target and response information across the two task demands, flexibly using shared coding schemes. Moreover, we evidenced emphasized neural representations of the instructions’ content under implementation demands. Altogether, these findings help to further clarify the neural mechanisms underlying task preparation in novel scenarios and represent a step forward to better characterize the proceduralization process.

## Funding

This work was supported by the Spanish Ministry of Science and Innovation (PID2019-111187GB-I00 to M.R., and IJC2019-040208-I to C.G.G), the European Union’s Horizon 2020 research and innovation program under the Marie Sklodowska-Curie (Ref. 835767, to C.G.G.), and the Spanish Ministry of Education, Culture and Sports (FPU17/01627 to A.S.)

## Acknowledgments

This research was part of A.S.’s activities for the Psychology Graduate Program of the University of Granada.

## CRediT author statement

Alberto Sobrado: Methodology, Formal analysis, Investigation, Visualization, Writing - original draft, Writing - review & editing.

Ana F. Palenciano: Conceptualization, Methodology, Formal analysis, Writing - review & editing, Software, Resources.

Carlos Gonzalez-García: Conceptualization, Methodology, Writing - review & editing.

María Ruz: Conceptualization, Writing - original draft, Writing - review & editing, Supervision, Project administration, Funding acquisition.

## Declaration of Competing Interest

The authors declare no competing financial interests.

## SUPPLEMENTARY MATERIAL

**Supplementary Figure 1.**
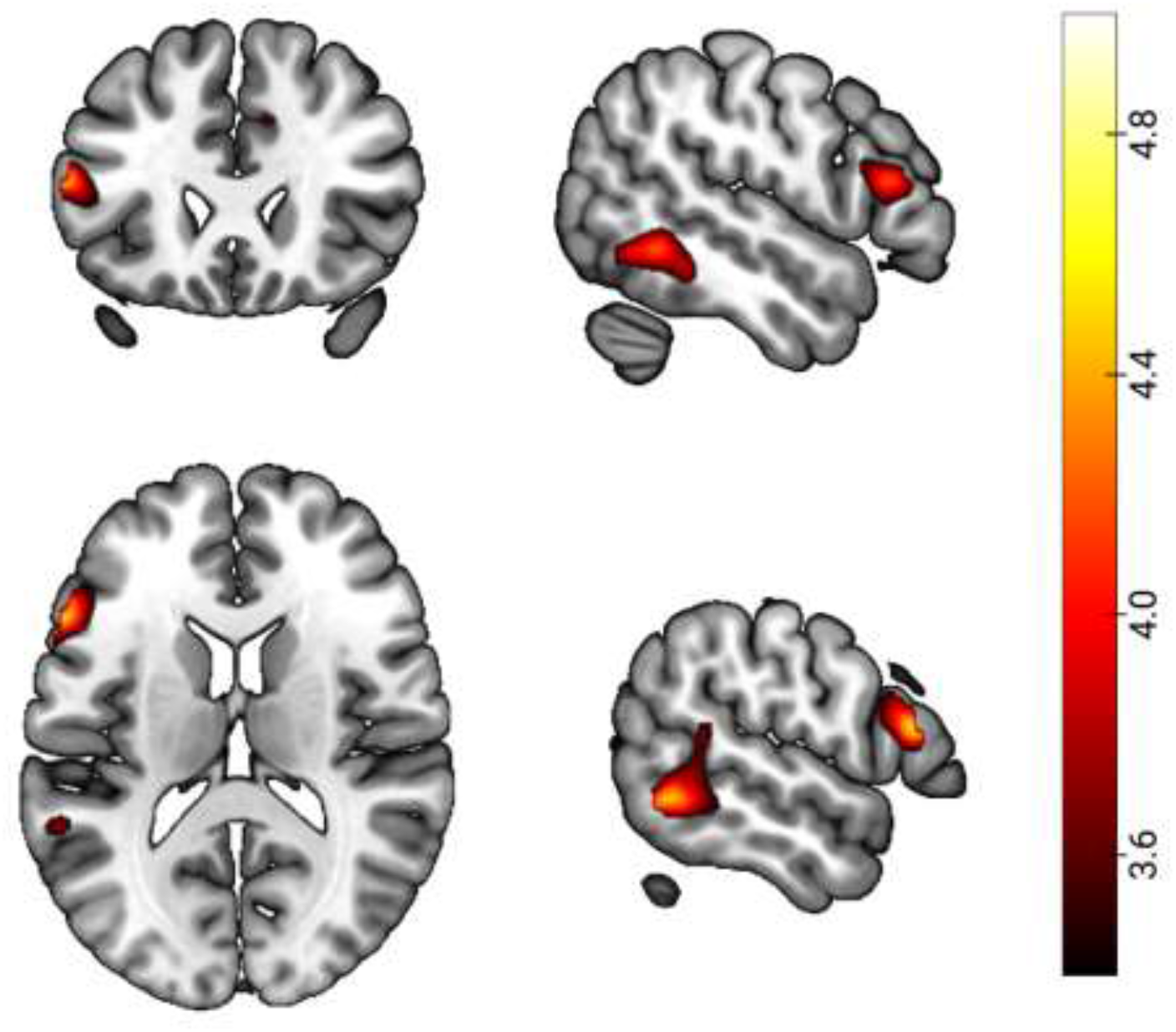
Uncorrected results (*p* < .001) from the whole-brain general decoding analysis with the dimension integration variable. The color shade indicates the *t*-values.

**Supplementary Figure 2.**
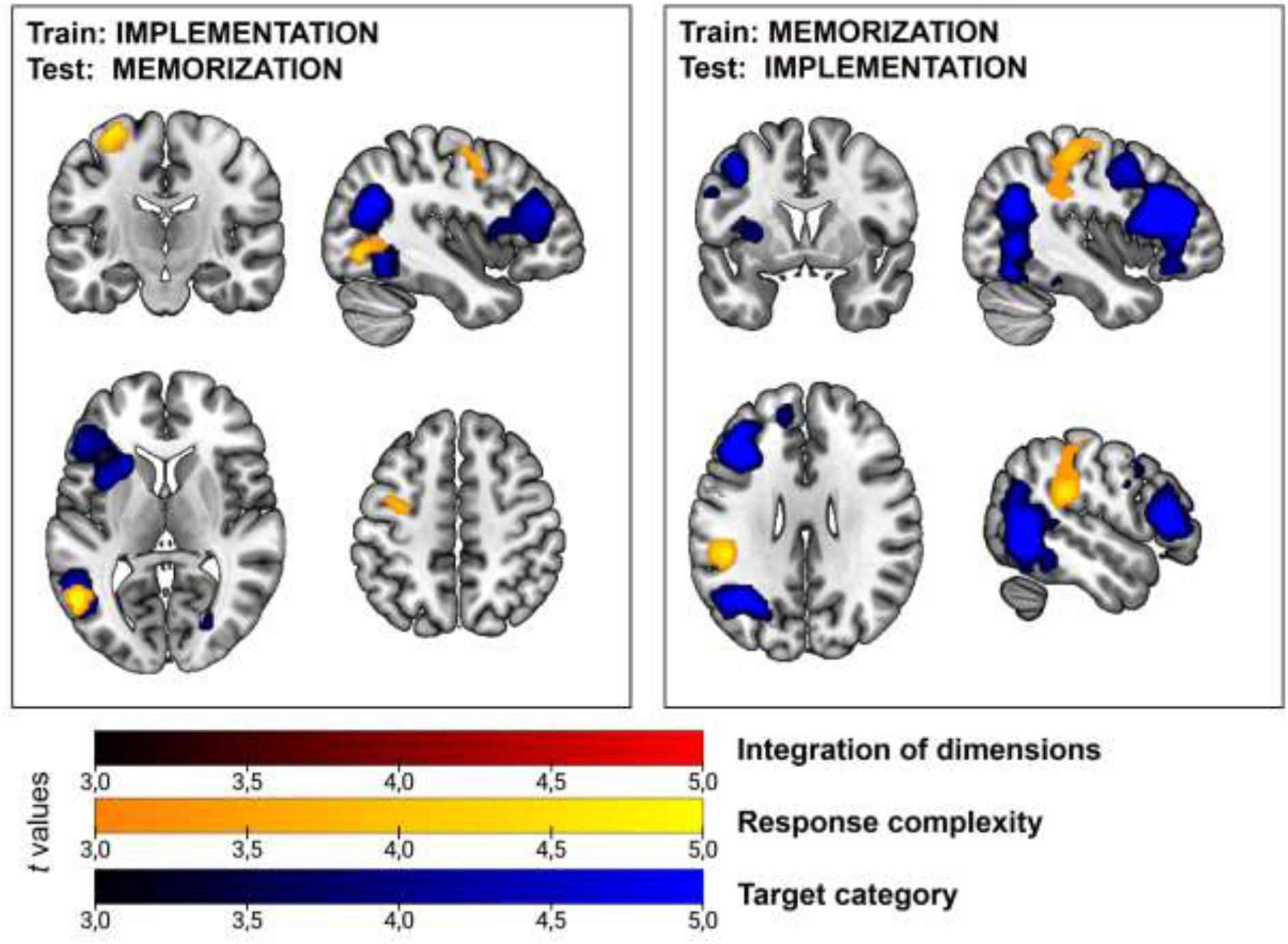
Results from the whole-brain cross-classification analysis, separately for each cross-classification direction (training on implementation and testing on memorization at the left, and the other way around at the right). Color shades indicate *t*-values (*p* < .05, FWE corrected for multiple comparisons).

**Supplementary Table 1.** Results of the behavioral repeated-measures ANOVAs. Only main effects and interaction terms of interest (i.e., two-way interactions between task demands and the remaining instruction’ variables) are shown.

| | *F* | *p* | $\eta_p^2$ |
| --- | --- | --- | --- |
| *Accuracy rates* |  |  |  |
| Task demands | 57.20 | <.001 | .65 |
| Dimension integration | 16.98 | <.001 | .35 |
| Response set complexity | 9.52 | .004 | .24 |
| Target category | 14.91 | <.001 | .33 |
| Task demands * Dimension integration | < 1 | .926 | <.01 |
| Task demands * Response set complexity | < 1 | .913 | <.01 |
| Task demands * Target category | 3.01 | .091 | .09 |
| *RT* |  |  |  |
| Task demands | 491.20 | <.001 | .94 |
| Dimension integration | 101.80 | <.001 | .77 |
| Response set complexity | 9.52 | .004 | .24 |
| Target category | .44 | .514 | .01 |
| Task demands * Dimension integration | 36.23 | <.001 | .54 |
| Task demands * Response set complexity | 17.02 | <.001 | .35 |
| Task demands * Target category | 29.70 | <.001 | .49 |

## Notes

### Competing Interest Statement

The authors have declared no competing interest.

